# Potassium channel-driven bioelectric signaling regulates metastasis in triple-negative breast cancer

**DOI:** 10.1101/2021.04.06.438714

**Authors:** Samantha L Payne, Priyanka Ram, Deepti H. Srinivasan, Thanh T. Le, Michael Levin, Madeleine J Oudin

## Abstract

There is a critical need to better understand the mechanisms that drive local cell invasion and metastasis to develop new therapeutics targeting metastatic disease. Bioelectricity is an important mediator of cellular processes and changes in the resting membrane potential (RMP) are associated with increased cancer cell invasion. However, the mechanism is not well understood. Our data demonstrate that altering the RMP of triple-negative breast cancer (TNBC) cells by manipulating potassium channel expression increases *in vitro* invasion, *in vivo* tumor growth, and metastasis, and is accompanied by changes in gene expression associated with cell adhesion. We describe a novel mechanism for RMP-mediated cell migration involving cadherin-11 and the MAPK pathway. Importantly, we identify a new strategy to target metastatic TNBC *in vivo* by repurposing FDA-approved potassium channel blockers. Our results provide an understanding of the mechanisms by which bioelectricity regulates cancer cell invasion and metastasis that could lead to a new class of therapeutics for patients with metastatic disease.

## Introduction

Despite identifying pro-metastatic cues and signaling pathways that drive invasion, a cellular process that supports tumor growth, dissemination, colonization, and ultimately metastasis, there are no clinically available treatments that directly target invading cells in cancer. As a result, metastasis remains the leading cause of death in cancer patients. The triple-negative breast cancer (TNBC) subtype accounts for approximately 15% of all breast cancer cases, and is associated with a poorer 5-year prognosis, increased likelihood of metastasis and shorter overall survival than the other subtypes^1,2^. Several clinical trials are ongoing to identify new agents to treat TNBC, however current strategies benefit only a subset of patients and have limited efficacy. Therefore, there is an urgent need for more effective treatments. Targeting bioelectricity-mediated signaling pathways of tumor cells is an emerging strategy to treat cancer^3–7^.

All cells generate and receive bioelectric signals through the flow of ions through transmembrane ion channel, transporters, and pumps, altering the resting membrane potential (RMP). Cellular RMP, the membrane potential at which the net ionic currents across the plasma membrane are zero, is maintained at a negative value^8^. The RMP is dependent on the concentration gradient of the most abundant ions (K^+^, Na^+^, Ca^2+^, and Cl^−^), membrane permeability to these ions, and ion channel expression and activity^8^. As result of changes in these properties, the RMP of cells can either become more negative (hyperpolarized) or more positive (depolarized). The bioelectric properties of cancer cells differ vastly from their normal counterparts: they are more depolarized and have dysregulated ion channel expression and activity^9–11^. Previous work has demonstrated that Na^+^ and K^+^ ion channel activity can regulate migration, invasion, and metastasis in breast cancer^12–15^. K^+^ channels, which selectively conduct K^+^ ions across the membrane down their electrochemical gradient, generally dominate the ion conductance at the resting state of normal cells, causing the RMP to be more positive than the equilibrium potential for potassium^8,16,17^. K^+^ channels can drive cell migration via modulation of intracellular calcium^14,18,19^, β-catenin signaling^20^ and changes in cell volume^21^. Furthermore, patients with apocrine breast cysts, which have a higher risk of developing into cancer, have an elevated K^+^ concentration in the cyst fluid^22^. Despite these observations, it remains unknown whether K^+^-driven cancer cell migration requires changes in the RMP and whether manipulating the RMP can be used to inhibit migrating cells. Ion channels are the second-most common target for existing pharmaceuticals and there are many already FDA-approved drugs targeting K^+^ channels available for non-cancer indications that could be repurposed for cancer therapy^23,24^. Understanding the role of the RMP and K^+^ channel activity in cell migration and cancer metastasis could lead to new treatment strategies for TNBC.

Here, we investigate the effect of manipulating the RMP through K^+^ channel-driven mechanisms on TNBC cell invasion and metastasis. We find, unexpectedly, that hyperpolarizing the RMP through overexpression of K^+^ ion channels increases cell migration, invasion, tumor growth and metastasis. For the first time, we characterize gene expression changes driven by hyperpolarization in TNBC cells, which lead to upregulation of pathways involved in cell adhesion, in particular the expression of cadherin-11. We demonstrate that cadherin-11-mediated MAPK signaling is involved in hyperpolarization-driven cell migration. Furthermore, blocking endogenous K^+^ channels in TNBC cells with the FDA-approved drug amiodarone causes RMP depolarization, decreasing *in vitro* migration and metastasis *in vivo*. Our results demonstrate that targeting K^+^ channel-mediated changes in RMP may provide a new strategy for treating metastatic disease.

## Results

### TNBC cell RMP is controlled by K^+^ ions

Given that the cellular RMP is determined by ion channel expression and activity, we first aimed to determine the ion channel expression profile of human breast cancers. Analysis of the TCGA dataset^23^ for gene expression of ion channel types revealed that K^+^ channel encoding genes are upregulated in invasive ductal carcinoma patient samples compared to normal breast tissue, whereas Na^+^ and Cl^−^ channel genes were both upregulated and downregulated (Figure 1a). Although it is known that K^+^ channel activity regulates the RMP of normal cell types, this has not been established for TNBC cells. To determine which ion species are dominant in setting the RMP of TNBC cells, we incubated human breast cancer cell lines (LM2, MDA-MB-231, SUM159, MDA-MB-468, and BT20) and immortalized normal breast epithelial MCF10A cells in a series of extracellular solutions that are balanced osmotically and differ only in the concentration of one ion^25^ (Figure S1). This allowed us to distinguish the influence of one ion species rather than targeting a single ion channel, by measuring the effect on the cellular RMP using DiBAC, a voltage-sensitive membrane dye^25,26^. We observed that incubating cells with increasing extracellular concentration of K^+^ (5.4mM to 135.4mM) results in significant depolarization of the cell population in metastatic lines LM2, MDA-MB-231, SUM159, and MDA-MB-468 (Figure 1b-e) but had no effect on the poorly metastatic BT20 or healthy epithelial MCF10A line (Figure 1f,g). Incubation in Na^+^ (28mM to 140mM) or Cl^−^ solutions (30.2mM to 151mM) had no effect on the RMP for any lines. Interestingly the increase in RMP was also associated with the reported metastatic potential of the cell lines^27^; in the presence of increased extracellular K^+^ concentration, lines with a higher metastatic potential depolarized more than low metastatic potential or healthy MCF10As. These data show that K+ channels are upregulated in human breast cancer tumors and suggest that TNBC cell RMP is regulated by the activity of K^+^ channels rather than that of Na^+^ or Cl^−^.

**Figure 1:**
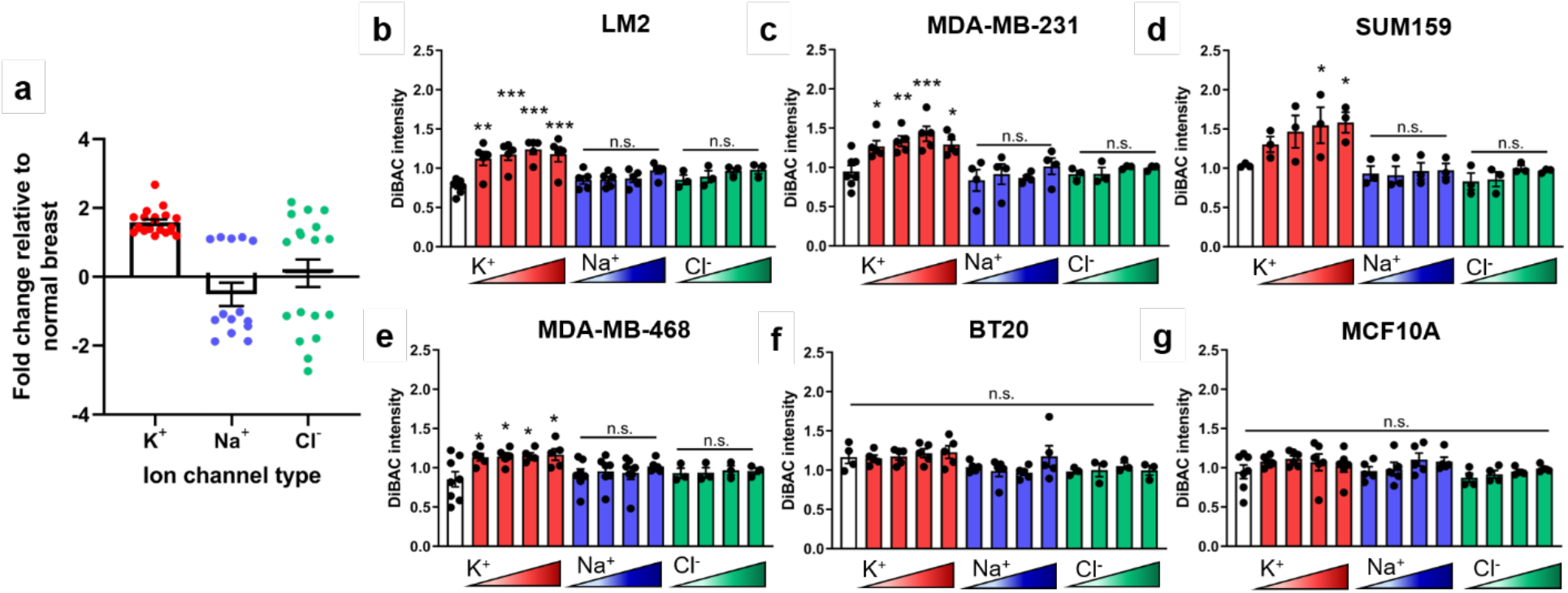
The RMP of metastatic TNBC cells is driven by K^+^ ions. **(a)** Fold change in ion channel mRNA levels in invasive ductal breast carcinoma relative to normal breast from TCGA dataset. The RMP of five breast cancer lines, **(b)** LM2 (**p=0.0004, ***p<0.0001), **(c)** MDA-MB-231 (*p=0.0415, **p=0.0084, ***p=0.0007), **(d)** SUM159 (*p=0.0489, 0.0331), **(e)** MDA-MB-468 (*p=0.0387, 0.0317, 0.0217, 0.0153), and **(f)** BT20, and one healthy epithelial breast cell line, **(g)** MCF10A, when incubated in ionic solutions of increasing concentration of a single ion, K^+^, Na^+^, or Cl^−^ and compared to physiological concentrations (white bar). Data are shown as mean ± S.E.M, n = 3-7 biological replicates. Significance was determined using a one-way ANOVA with Dunnett’s post hoc test to compare each set of ionic solutions to physiological concentration.

### K^+^ channel-driven RMP hyperpolarization increases TNBC cell invasion

To study the effect of K^+^ conductance and change in the RMP on breast cancer cell metastatic potential, we engineered MDA-MB-231 and MDA-MB-468 cell lines to stably express two different types of GFP-tagged K^+^ channels: Kv1.5, a voltage-gated channel (231-Kv1.5 and 468-Kv1.5), and Kir2.1, a constitutively open, inwardly-rectifying channel (231-Kir2.1 and 468-Kir2.1) (Figure 2a). These channels were chosen as a tool to hyperpolarize the RMP, as has been previously demonstrated^28–31^ and compared against their respective negative control expressing a fluorophore alone. The observed nuclear localization of GFP in the Kv1.5 expressing cells (Figure 2a) is due to the *Thosea asigna* virus 2A (T2A) sequence linking Kv1.5 and histone GFP in the Kv1.5 construct, which during translation initiates ribosomal skipping and as a result the histone GFP is cleaved and shuttled to the nucleus^32^. Expression of either channel resulted in a significantly hyperpolarized RMP in both cell lines, with a change in voltage of approximately −18mV for Kv1.5 and −40mV for Kir2.1 (Figure 2b,c). We confirmed that this change in RMP was accompanied by an increase in outward K^+^ current for both MDA-MB-231 and MDA-MB-468 derived lines (Figure S2a,c), although the amount of current measured varied considerably between the two lines, with both 231-Kv1.5 and - Kir2.1 lines generating a higher K^+^ current than their MDA-MB-468 counterparts. We also confirmed that pharmacological blocking of the channels with an inhibitor specific to each channel type resulted in a reversal of the observed hyperpolarization (Figure S2b,d).

**Figure 2:**
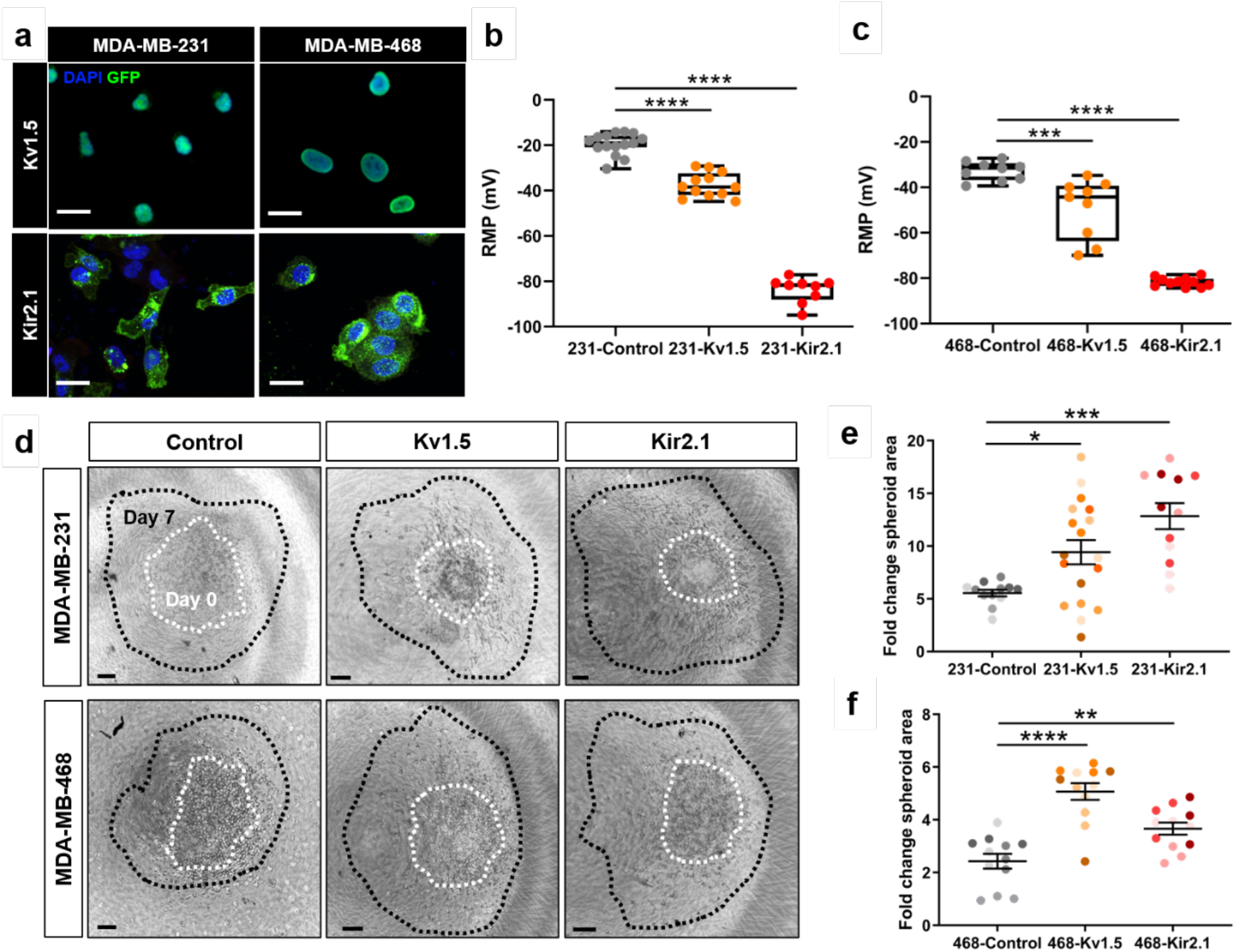
Overexpression of K^+^ channels induces RMP hyperpolarization and increases 3D cell invasion. **(a)**Expression of GFP-tagged Kv1.5 and Kir2.1 channel constructs by lentiviral transduction into MDA-MB-231 and MDA-MB-468 breast cancer cell lines visualized with GFP. Scale bar = 10μm. Electrophysiology measurement of the RMP in K^+^ channel overexpressing lines compared to negative controls in **(b)** MDA-MB-231 (****p<0.0001) and **(c)** MDA-MB-468 (***p=0.0002, ****p<0.0001) cells. **(d)** Representative images at day 7 (black) overlayed with outlines of spheroid area at day 0 (white). Scale bar = 100μm. The fold change in spheroid area in **(e)** MDA-MB-231 (*p=0.0246, ***p=0.0001) and **(f)** MDA-MB-468 (**p=0.0066, ****p<0.0001) lines. Data are pooled from three or more biological replicates and shown as mean ± S.E.M. Different color shades within group represent samples from different replicates. Significance was determined using a one-way ANOVA with Dunnett’s multiple comparisons test.

Next, we investigated the impact of K^+^ channel-driven hyperpolarization on 3D cell invasion. Spheroids made from control and K^+^ channel-expressing cells were encapsulated in collagen I matrices and grown for seven days. The change in spheroid area was quantified as a measurement of cell invasion into the surrounding matrix, a model which most accurately reflects the physical properties of tumors^33^. We found that overexpression of Kv1.5 and Kir2.1 significantly increased 3D cell invasion in both cell lines when compared to the negative control (Figure 2d-f). In addition, we evaluated the effects of the channels on 2D migration, demonstrating an increase in cell speed in the 231-Kv1.5 and 231-Kir2.1 lines as compared to the control (Figure S3a-e). Lastly, cell proliferation was measured by quantification of Ki67 expression in cells. In the MDA-MB-231 lines, Kv1.5 overexpression had no effect on proliferation, whereas in the MDA-MB-231-Kir2.1 and MDA-MB-468 lines proliferation was decreased compared to the negative controls (Figure S3f,g). Taken together, these data demonstrate that K^+^ channel-driven RMP hyperpolarization increases TNBC cell migration and invasion.

We next investigated the effect of K^+^ channel-driven RMP hyperpolarization on *in vivo* tumor growth and metastasis in a mouse xenograft model. NOD-SCID-γ mice were injected with 231-Kv1.5, 231-Kir2.1, or 231-Control lines in the 4^th^ mammary fat pad and tumor growth was tracked over time. From four to seven weeks post-injection, tumor volume was significantly greater in mice in the 231-Kv1.5 and 231-Kir2.1 group compared to the 231-Control group (Figure 3a). Animals were sacrificed at seven weeks and metastasis to the lungs was quantified using H&E staining and expressed as the number of metastases relative to total lung area (Figure 3b) and the number of metastases relative to final tumor volume (metastatic index, Figure 3c). Our data show that both the 231-Kv1.5 and 231-Kir2.1 group had significantly more metastases present than the 231-Control group (Figure 3d-f). Combined, these data demonstrate that K^+^ channel-driven RMP hyperpolarization enhances TNBC cell migration and invasion *in vitro*, and tumor growth and metastasis *in vivo*.

**Figure 3:**
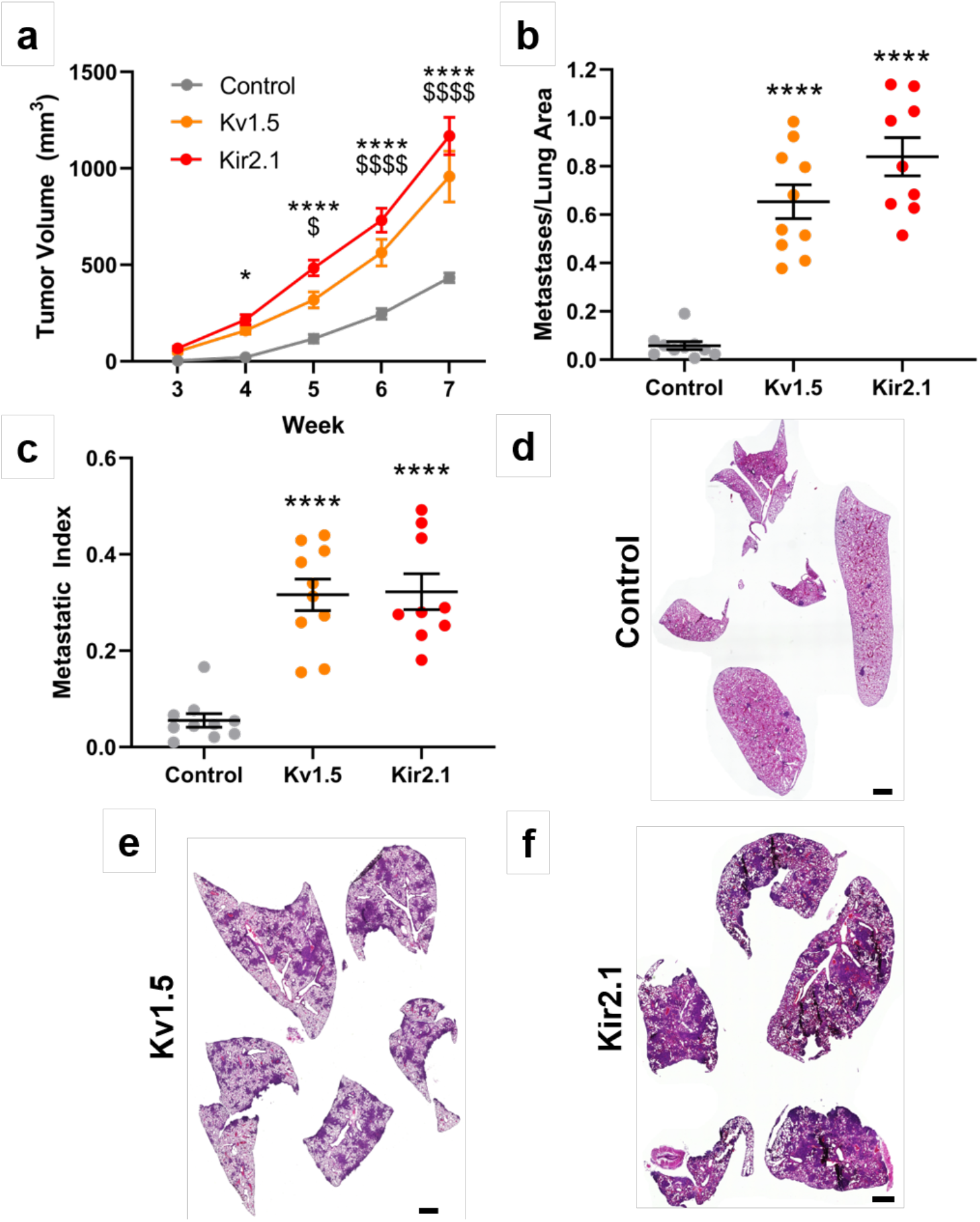
K^+^ channel-driven RMP hyperpolarization increases TNBC tumor growth and metastasis. **(a)** Tumor volume measured over time in 231-Kv1.5 and 231-Kir2.1 volume compared to the 231-Control group. *p=0.0191, ****p<0.0001 for 231-Kir2.1 compared to 231-Control; $p=0.0153, $$$$p<0.0001 for 231-Kv1.5 compared to 231-Control. **(b)** The number of metastases per lung and **(c)** the metastatic index was significantly increased in the 231-Kv1.5 and Kir2.1 group compared to 231-Control (****p<0.0001). Significance was determined using a one-way ANOVA with Dunnett’s multiple comparisons test. Representative lung tissue sections stained with H&E for **(d)** 231-Control, **(e)** 231-Kv1.5, and **(f)** 231-Kir2.1 group. Scale bar = 1mm. Data are shown as mean ± S.E.M. N = 9-10 animals/group.

### K^+^ channel-driven RMP hyperpolarization drives upregulation of genes associated with cell adhesion and MAPK signaling

After determining that overexpression of K^+^ channels and hyperpolarization result in profound phenotypic changes in TNBC cell lines, we aimed to characterize the changes in gene expression associated with TNBC cell hyperpolarization by performing RNA sequencing on the 231-Kv1.5, 231-Kir2.1 and 231-Control lines. Differential expression analysis revealed a significant number of genes to be differentially expressed in channel overexpressing lines compared to the control (Figure 4a). Principal component (PC) analysis determined that 75% of the overall variance was found in PC1 between groups separating control and K^+^ channel-expressing cells, indicating that changes in the cellular RMP have profound effects on gene expression (Figure 4b). To focus on genes that were commonly dysregulated in both 231-Kv1.5 and 231-Kir2.1, we combined differentially-expressed genes from both lines and analyzed the average fold change in their expression compared to the 231-Control group, generating a list of significantly upregulated and downregulated genes (Figure 4c). We next analyzed enrichment for functional pathways and generated normalized enrichment scores for differentially expressed genes shared by both 231-Kv1.5 and 231-Kir2.1 lines, using a false discovery rate (FDR) threshold of 0.1 and significance of p<0.05. Relevant positively enriched pathways include ECM regulators, ERK1/2 regulation, MAPK activity, cell adhesion and cell migration (Figure 4d). Together, these results demonstrate that K^+^ channel-driven RMP hyperpolarization drives significant changes in gene expression, upregulation of pathways involved in cell adhesion.

**Figure 4:**
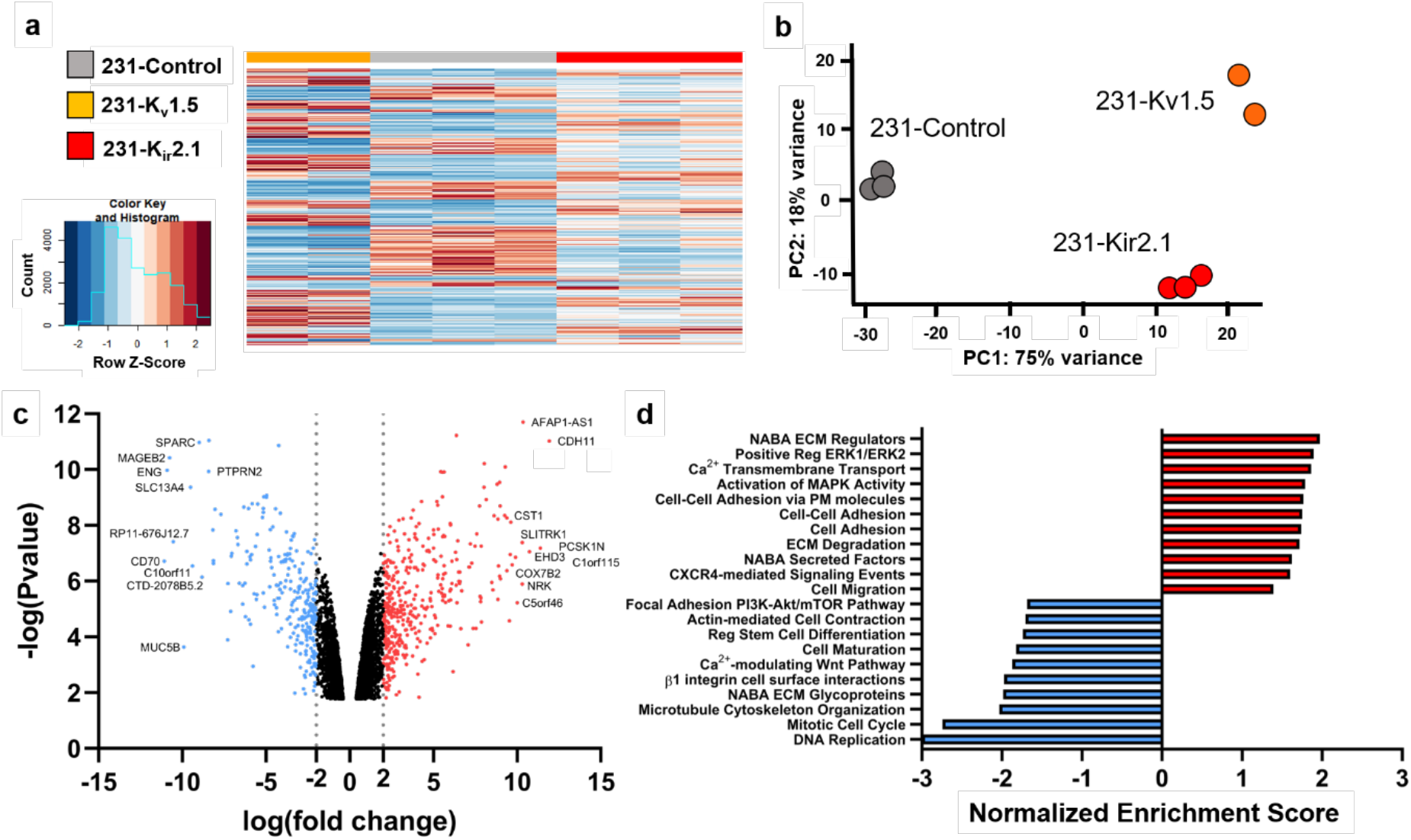
K^+^ channel-driven RMP hyperpolarization upregulates genes associated with cell adhesion. **(a)** Heat map clustering of differentially expressed genes in 231-Kv1.5 and 231-Kir2.1 compared to 231-Control. **(b)** Principal component (PC) analysis showing clustering and variance captured for cell lines. **(c)** Volcano plot demonstrating log(fold change) and log(p value) of upregulated and downregulated gene expression in the combined group of 231-Kv1.5 and 231-Kir2.1. The top 10 genes with highest fold change are labeled. **(d)** Normalized enrichment score for a subset of functional pathways upregulated and downregulated in the combined 231-Kv1.5/231-Kir2.1 group compared to 231-Control.

The observed upregulation of genes involved in cell adhesion suggests that overexpression of K^+^ channels may alter the adhesion and morphology of TNBC cells, which our lab and others have demonstrated is highly predictive of 3D invasive behavior driving metastasis^34,35^. We characterized cell area and morphology of MDA-MB-231 K^+^ overexpressing lines cultured on collagen I-coated glass plates. Quantification of cell area using F-actin staining (Figure 5a) revealed that 231-Kv1.5 and 231-Kir2.1 have a significantly larger area than 231-Control cells (Figure 5b). Cell compactness, the ratio of the area of a cell compared to a circle of the same perimeter, and eccentricity, the ratio of the minor to major cell axis, were found to be increased in 231-Kir2.1 cells compared to 231-Control and 231-Kv1.5 cells (Figure 5c, d). Cell solidity, the ratio of the area of a cell to a convex hull of the cell, was significantly decreased (Figure 5e). These morphological features indicate that 231-Kir2.1 cells are more elongated, have more protrusions, and greater boundary irregularity compared to 231-Control and 231-Kv1.5 cells. An important step in cell migration is the formation of focal adhesions between the intracellular actin cytoskeleton and the ECM. To test if K^+^ channel-driven hyperpolarization is linked to focal adhesion formation, we investigated levels of phosphorylated focal adhesion kinase protein (pFAK397), a regulator of adhesion turnover, and phosphorylated paxillin (pPax118), a focal adhesion adaptor protein phosphorylated by FAK, in our channel overexpressing cell lines. We found that 231-Kv1.5 and 231-Kir2.1 cells have significantly higher levels of pFAK when compared to 231-Control cells (Figure 5f,g), but did not detect a difference in pPax118 expression (Figure 5h). Combined, our results suggest that K^+^ channel-driven RMP hyperpolarization is associated with changes in cell shape and focal adhesion signaling.

**Figure 5:**
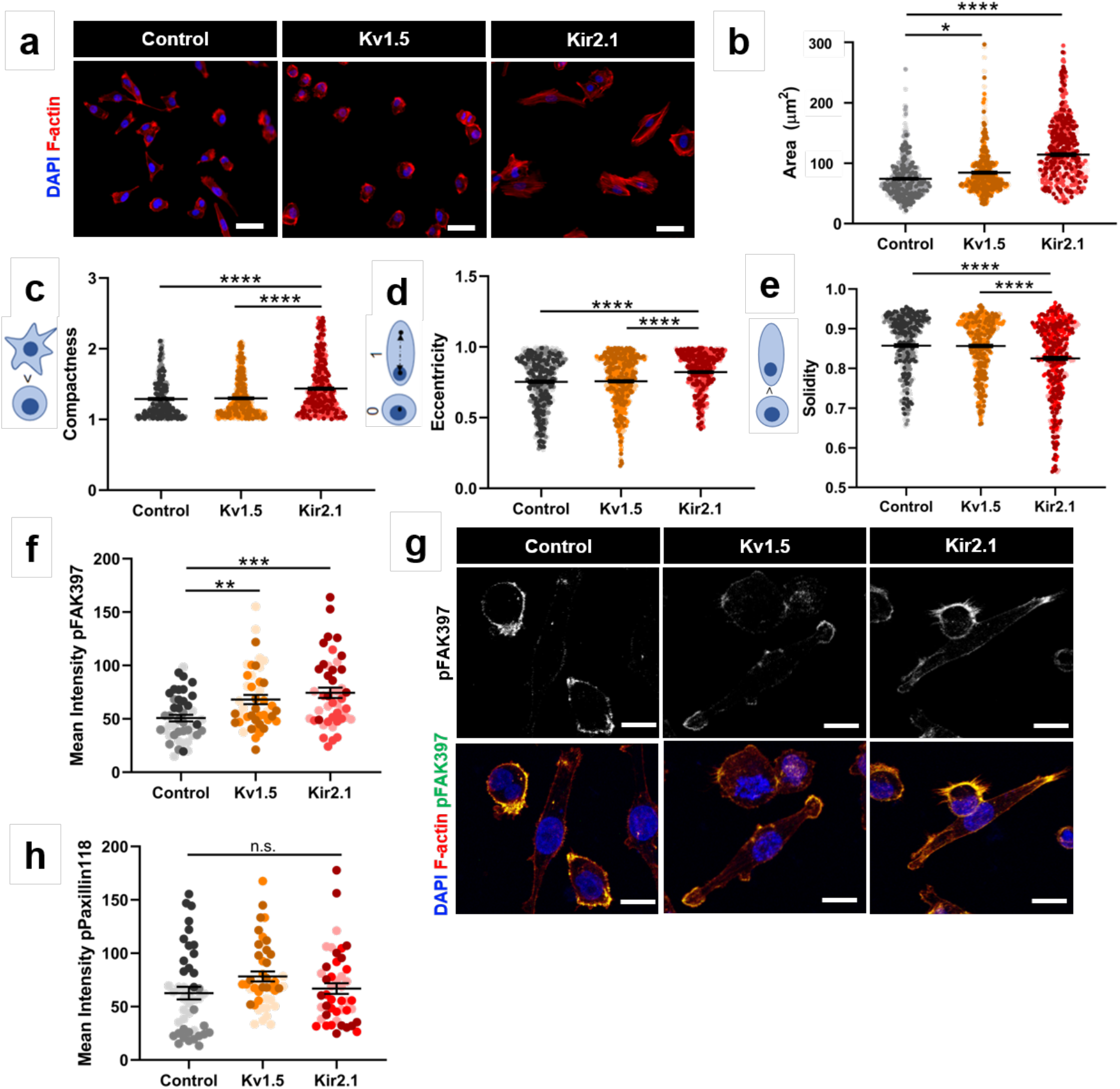
K^+^ channel-driven RMP hyperpolarization alters cell morphology and focal adhesion signaling. **(a)** Representative images of overexpressing cell lines stained with phalloidin for F-actin. Scale bar = 10μm. **(b)** Quantification of F-actin-based cell area (*p=0.0202, ****p<0.0001). Quantification of **(c)** compactness, **(d)** eccentricity, and **(f)** solidity (****p<0.0001). **(f)** Quantification of pFAK397 expression in overexpressing lines compared to 231-Control (**p=0.0065, ***p=0.0002). **(g)** Representative images of pFAK397 expression. Scale bar = 10μm. **(h)** Quantification of pPax118 in overexpressing lines compared to 231-Control. Data are pooled from three or more biological replicates and shown as mean ± S.E.M. Different color shades within group represent samples from different replicates. Significance was determined using a one-way ANOVA with Dunnett’s multiple comparisons test.

### Cadherin-11 is necessary for K^+^ channel-driven cell migration

Cadherin-11 is a member of the cadherin super-family which mediates homophilic cell-cell adhesion in a calcium-dependent manner by interacting with the cytoplasmatic catenins, α, β, and γ, and has been shown to drive metastasis in epithelial cancers^36–38^. Due to its significant 11-fold genetic upregulation in the 231-Kv1.5 and 231-Kir2.1 cell lines, we chose to investigate the role of cadherin-11 in the K^+^ channel-driven effects observed. To determine if the increased expression of the *CDH11* gene seen in the RNAseq (Figure 4c) corresponds to increased cadherin-11 protein expression, we quantified cadherin-11 in K^+^-overexpressing 231 cell lines. Compared to 231-Control cells, 231-Kv1.5 and 231-Kir2.1 expressed significantly higher levels of cadherin-11 as determined by immunocytochemistry (Figure 6a,b) and western blot (Figure 6c,d). We next knocked down expression of *CDH11* using siRNA which has been used previously in breast cancer cells^41^ and measured the effects on cell migration and adhesion. An 80-90% reduction in *CDH11* expression was achieved with siRNA knockdown after 48 hours which was verified with cadherin-11 immunostaining (Figure S4a) and western blot (Figure S4b,c). This *CDH11* knockdown resulted in a significant decrease in 2D cell migration speed in both 231-Kv1.5 and 231-Kir2.1 cell lines (Figure 6e) as well as a decrease in cell area (Figure 6f). Cadherin-11 has been previously linked to ERK phosphorylation^38,39^, and ERK in turn can promote cell migration by phosphorylating FAK and Paxillin to drive focal adhesion formation^40^. Furthermore, MAPK signaling pathway-related genes were enriched in our RNAseq dataset (Figure 4d). Consistent with this, we observed that knockdown of *CDH11* results in a significant decrease in the ratio of pERK1/2 to total ERK1/2 (Figure 6g,h). These results suggest that the increase in 2D migration of hyperpolarized cells is driven by cadherin-11 and MAPK signaling.

**Figure 6:**
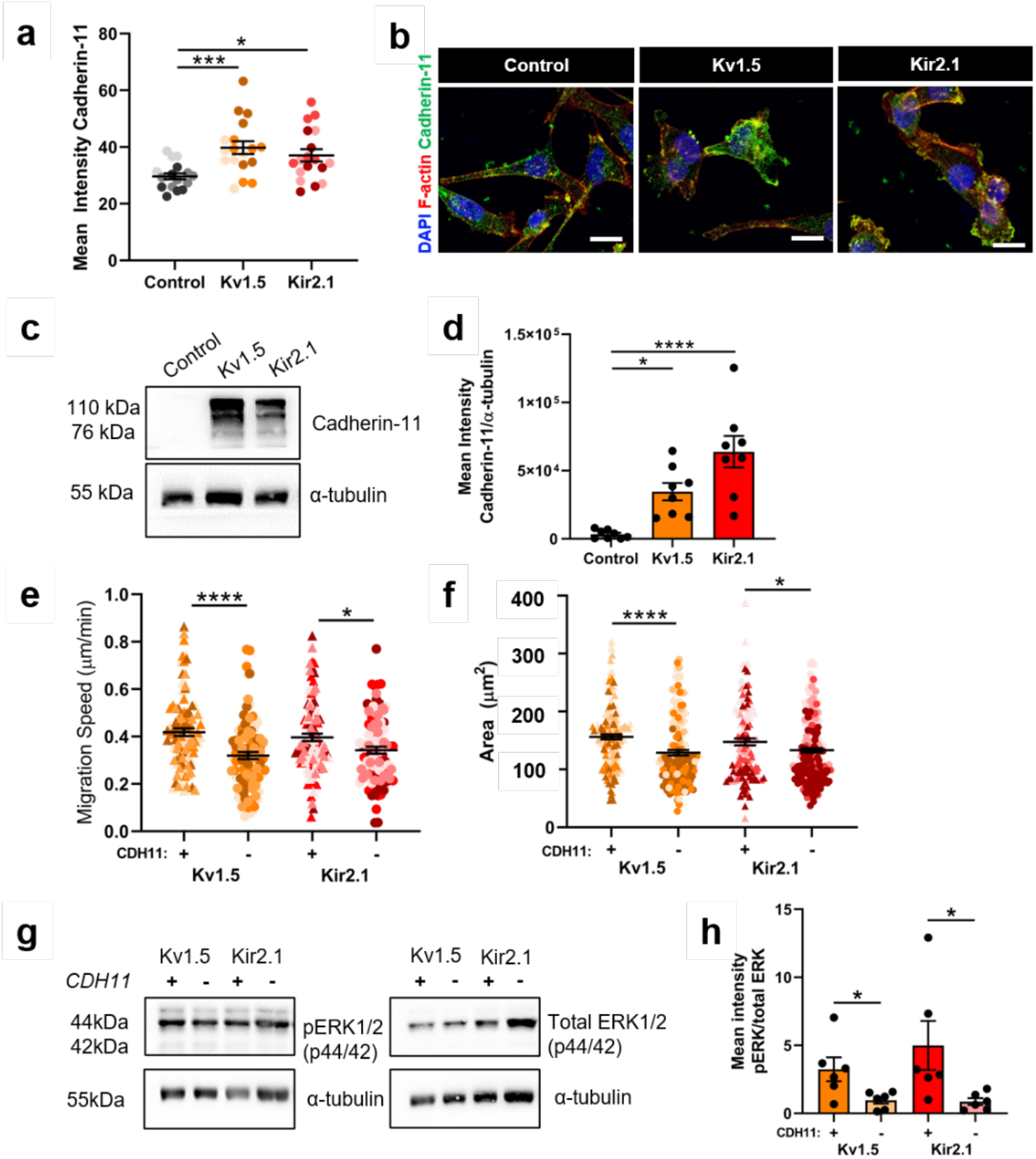
Cadherin-11 mediates the pro-migratory effects of K^+^ channel-driven hyperpolarization. **(a)** Quantification of cadherin-11 expression in 231 cell lines (*p=0.0168, ****p<0.0001). **(b)** Representative images of cadherin-11 expression (green) colocalized with F-actin (red). Scale bar = 10μm. **(c)** Representative western blot and **(d)** quantification of cadherin-11 expression in 231 cell lines (*p=0.0498, ***p=0.0007). **(e)** 2D migration speed following *CDH11* knockdown by siRNA relative to control siRNA (*p=0.0143, ****p<0.0001). **(f)** Quantification of F-actin-based cell area following *CDH11* knockdown by siRNA relative to control siRNA (*p=0.0363, ****p<0.0001). **(g)** Representative western blot and **(h)** quantification of the ratio of pERK1/2 to total ERK1/2 with and without *CDH11* knockdown (*p=0.032 for Kv1.5, *p=0.0471 for Kir2.1). Data are pooled from three or more biological replicates and shown as mean ± S.E.M. Different color shades within group represent samples from different replicates. Significance was determined using a one-way ANOVA with Dunnett’s multiple comparisons test.

### Pharmacological inhibition of K^+^ channels decreases cell migration

Our results using K^+^ channel overexpressing breast cancer lines suggests that K^+^ channel activity and the subsequent hyperpolarization of the RMP can increase cell invasion and metastasis. To determine the feasibility of targeting the RMP of breast cancer cells to treat metastatic disease, we next investigated the effect of K^+^ channel blockers on cell invasion and metastasis. We mined the online repurposed drug database ReDO^23,42^ and selected four clinically available drugs that are known to block K^+^ channels and have been associated with anti-cancer effects in other tumor types^42^: amiodarone, carvedilol, imipramine, and thioridazine. An initial *in vitro* screen was performed with MDA-MB-231 and MDA-MB-468 cell lines, with all drugs inducing significant tumor cell death with IC50s in the low micromolar range (Figure 7a; Figure S5a-c). Next, we measured the change in RMP elicited by each drug using DiBAC at two doses in the range of the IC50 and determined that all drugs tested caused depolarization of the RMP in MDA-MB-231 and MDA-MB-468 cells (Figure 7b-e; Figure S5d,e). Due to the relatively large depolarization produced by amiodarone, we moved forward with this drug for subsequent experiments. Amiodarone is a class III anti-arrhythmic drug known to block voltage-gated and inward rectifier K^+^ channels^43–45^ and has been shown improve survival of hepatocellular carcinoma patients^46^. Treatment of MDA-MB-231 and MDA-MB-468 cells with amiodarone at 5 and 10μM resulted in a significant reduction in 2D cell migration for MDA-MB-231 cells (Figure 7f). We also saw a decrease in proliferation of MDA-MB-231 and MDA-MB-468 cells treated with 10μM amiodarone as measured by Ki67 expression (Figure S5f,g). Interestingly, we also observed a reduction in cadherin-11 expression with amiodarone treatment of approximately 70% and 50% with 5μM and 10μM respectively (Figure 7g). Lastly, we investigated the effect of blocking K^+^ channels with amiodarone on *in vivo* tumor growth and metastasis in a mouse xenograft model. NOD-SCID-γ mice were injected with MDA-MB-231 cells in the 4^th^ mammary fat pad and tumor growth was tracked over time. When tumors reached an average volume of 200mm^3^ mice received daily i.p. injections of 0.05mg/kg amiodarone or DMSO for 14 days, followed by 0.5mg/kg amiodarone or DMSO for 10 days. We observed a trend towards a lower tumor volume in animals treated with amiodarone compared to the DMSO control but it was not statistically significant (Figure S5h). Quantification of the metastases in the lungs at sacrifice revealed that animals treated with amiodarone had significantly fewer metastases than vehicle-treated animals (Figure 7h-j). Overall, our results demonstrate that depolarization of TNBC cells via amiodarone treatment leads to decreased migration and a reduction in the number of lung metastases *in vivo*.

**Figure 7:**
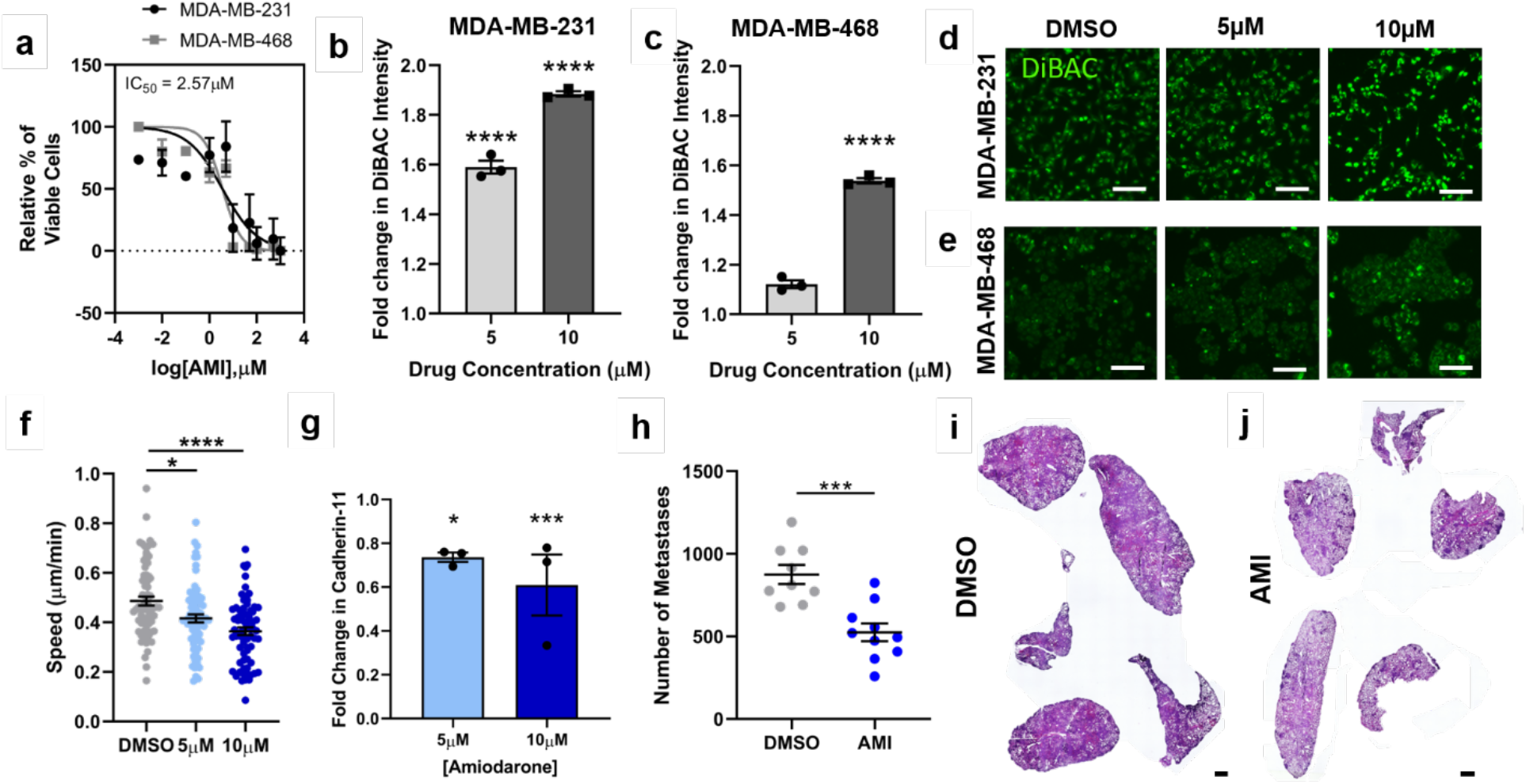
Amiodarone treatment depolarizes TNBC RMP resulting in decreased cell migration and lung metastases. **(a)** Dose response curves of cell viability in MDA-MB-231 and MDA-MB-468 cells cultured with varying concentrations of amiodarone (AMI). RMP of **(b)** MDA-MB-231 and **(c)** MDA-MB-468 cells measured by fold change in DiBAC intensity after amiodarone treatment at IC50 concentrations (****p<0.0001). Representative images of **(d)** MDA-MB-231 and **(e)** MDA-MB-468 cells treated with amiodarone or vehicle and stained with DiBAC. Scale bar = 60μm. **(f)** Cell migration speed following treatment with amiodarone in MDA-MB-231 cells (*p=0.0167, ****p<0.0001). Data are pooled from three or more biological replicates and shown as mean ± S.E.M. Significance was determined using a one-way ANOVA with Dunnett’s multiple comparisons test. **(g)** Fold change in cadherin-11 expression with amiodarone treatment in vitro quantified from western blot (*p=0.0486, ***p=0.0002). **(h)** Number of metastases per area of lung after daily i.p. injections of amiodarone at 0.05mg/kg (14 days) followed by at 0.5mg/kg (10 days) or DMSO for a total of 24 days (***p=0.0003). Representative lung tissue sections from animals treated with **(i)** DMSO or **(j)** amiodarone stained with H&E. Scale bar = 1mm. Significance was determined using a Student’s t test. N = 9-10 animals per group.

## Discussion

Here we show that hyperpolarization of TNBC cells through overexpression of K^+^ channels leads to enhanced cell invasion, tumor growth, and metastasis. We provide the first characterization of hyperpolarization-induced changes in gene expression in cancer, which included upregulation of genes associated with cell adhesion and MAPK signaling. Based on this, we also identify a novel hyperpolarization-driven mechanism of cell migration mediated by cadherin-11 and MAPK signaling. These effects were observed with the overexpression of two different types of K^+^ channels whose activity is modulated by different stimuli, suggesting that the effects seen are a result of the RMP hyperpolarization and not a channel-specific phenomenon. Lastly, we demonstrate that the bioelectric state of breast cancer cells can be targeted pharmacologically using the K^+^ channel blocker amiodarone to reduce breast cancer cell migration and lung metastasis in a mouse model of TNBC.

Previously it has been reported that breast cancer cells are less negative than their normal counterparts and that blocking sodium channels inhibits cell migration and invasion in MDA-MB-231 and MCF7 cells^9,47,48^, however if these effects are RMP-dependent is not noted. In contrast, our results suggest that hyperpolarization via K^+^ channel expression leads to increased breast cancer cell invasion. In line with our results are other reports that hyperpolarization can drive migration in cancer and other cell types^14,49,50^. For example, blocking K^+^ channel KCa2.3 activity in both hyperpolarized melanoma cells and MDA-MB-435 breast cancer cells inhibits 3D Matrigel invasion^49^ and 2D migration^51^, respectively. Furthermore, hyperpolarization has also been linked to a redistribution of F-actin towards the plasma membrane and an increase in adherens junction stability in bovine corneal endothelial cells^52^, consistent with our observed upregulation of cell adhesion-associated genes. Our results also support a model of hyperpolarization-driven transcription of *CDH11* and increased cadherin-11 levels in TNBC cells. Cadherin-11 has been previously shown to drive cell invasion and metastasis by two main pathways. First, it regulates TNBC cell migration by increasing β-catenin nuclear localization and the transcription of Wnt targets Met, c-Myc, c-Jun, MMP7, Sox2, CD44, KLF4^53^. However, the mRNA levels of these targets were unchanged in our dataset with the MDA-MB-231 or MDA-MB-468. Secondly, Cadherin-11 promotes paxillin phosphorylation, and adhesion formation^38^, as well as Rac-driven actin dynamics^40^, leading to downstream MAPK signaling and cell motility. Although further work will be needed to fully elucidate the mechanisms involved, overall, our results suggest that control of cancer cell migration and invasion by the RMP may be more complex than previously appreciated.

In addition to demonstrating that TNBC RMP is an important regulator of cell migration, invasion, and metastasis, we also show that it can be used to target metastasis by depolarizing MDA-MB-231 cells with amiodarone. It is worth noting that in addition to blocking outward potassium channels, amiodarone is reported to block inward sodium and calcium currents in cardiac cells under certain conditions^54–56^ therefore we cannot fully attribute the observed depolarization of MDA-MB-231 cells with amiodarone to potassium channel blocking alone. Nevertheless, we show that the depolarization produced by amiodarone treatment results in reduced TNBC cell migration and metastasis, demonstrating that the cellular RMP is a targetable feature of TNBC cells that can be exploited to treat metastasis. Interestingly, amiodarone has also been shown to enhance the *in vitro* toxicity of standard of care chemotherapy drugs for TNBC such as doxorubicin, albeit in other tumor types^57–59^. Given that chemotherapeutic drugs impact cell proliferation, and that transient hyperpolarization is needed for progression through the cell cycle^60–62^, it is possible that disruption of the cellular RMP by blocking K^+^ channels with amiodarone also could enhance the effect of chemotherapeutic drugs.

In conclusion, we demonstrate that TNBC cell hyperpolarization drives migration, invasion and metastasis which can be targeted by depolarizing the RMP pharmacologically. This hyperpolarization is associated with striking changes in gene expression including upregulation of cell adhesion and MAPK pathways involving cadherin-11 signaling. Our results reveal the importance of RMP for pro-migratory signaling pathways in cancer and identify a new therapeutic target for breast cancer metastasis.

## Methods

### Reagents

Reagents were purchased from Fisher Scientific (Hampton, NH) or Sigma (St. Louis, MO) unless otherwise specified. Antibodies used were: anti-α-Tubulin (T9026; Sigma, St. Louis, MO), anti-pFAK397 (3283; Cell Signaling Technology, Danvers, MA), anti-pPaxillin118 (44-722G; ThermoFisher Scientific, Waltham, MA), anti-cadherin-11 (71-7600; ThermoFisher Scientific, Waltham, MA), anti–phospho-p44/42 ERK (Thr202/Tyr204; 4370; Cell Signaling Technology, Danvers, MA), anti-p44/42 ERK (9102; Cell Signaling Technology, Danvers, MA), anti-Ki67 (ab15580; Abcam, Cambridge, UK), AlexaFluor 488 anti-rabbit secondary (A-11008; ThermoFisher Scientific, Waltham, MA), and AlexaFluor 647 anti-rabbit secondary (A-21244; ThermoFisher Scientific, Waltham, MA). ECM substrates used were: Collagen I (CB-40236; Fisher Scientific, Hampton, NH). Inhibitors used were: Amiodarone hydrochloride (40-955-0; Tocris Bioscience, Minneapolis, MN), Carvedilol (C3993,Sigma, St. Louis, MO), Imipramine hydrochloride (I7379; Sigma, St. Louis, MO), and Thioridazine hydrochloride (T9025, Sigma, St. Louis, MO).

### Cell lines

The MDA-MB-231 and MDA-MB-468 breast cancer cell lines, and the HEK293T cell line were obtained from ATCC (Manassa, VA) and cultured in DMEM with 10% FBS (SH30071.03; Cytiva; Marlborough, MA) and penicillin-streptomycin glutamine (10378016; ThermoFisher Scientific; Waltham, MA). Cells were passaged regularly when they reached approximately 70% confluency and used between p5 and p15 for all experiments. Cells were routinely checked for the presence of mycoplasma by a PCR based method using a Universal Mycoplasma Detection Kit (30-1012K; ATCC, Manassas, VA). Only mycoplasma negative cells were used in this study.

### Plasmids and lentiviral infection

Viruses were produced as described previously^31^. For infections, 1μg of constructs expressing human ion channel coding proteins RFP, Kv1.5-T2A-his-EGFP, or KCNJ2(Kir2.1)-Y242F-EGFP were used. For each construct, HEK293T cells at a confluency of 80% were used to transfect for virus production. Constructs were mixed with 0.5μg helper (encoding gag/pol) and 0.1μg encoding coat protein (pMD2) along with 6μL TransIT-Lenti transfection reagent (MIR6603; Mirus Bio; Madison, WI) in 200uL Opti-MEM (31985062; ThermoFisher Scientific; Waltham, MA). Culture media of HEK293T cells was replaced with Opti-MEM and 200μL of transfection mixture and cells were kept in transfection media overnight. Transfection media was replaced with DMEM containing 1% Pen/Strep the following day. Two days post-transfection media containing virus was collected and stored at −80C. Collection was repeated on the following day and the total virus collected was combined. To transduce cells, MDA-MB-231 or MDA-MB-468 cells were seeded in six-well plates at 2 × 10^5^ cells per well in 2mL DMEM containing 10μg/mL polybrene and virus. The plates were centrifuged at 1500 r.p.m., 4°C for 1 h and incubated at 37 °C. Medium was changed after 16 h. Infected cells were sorted by FACS and gating on eGFP-derived green fluorescence.

### Voltage sensitive dye

Changes in RMP were measured using DiBAC (ThermoFisher Scientific, Waltham, MA) as described previously^25^. MDA-MB-231 or MDA-MB-468 cells, cultured in black clear-bottomed 96-well plates, were rinsed once with 200μL of Fluorobrite^TM^ medium (ThermoFisher Scientific, Waltham, MA) and solutions containing 1 μM DiBAC were added to cells and incubated in a cell incubator for 15 min to ensure dye distribution across the membrane. Assays were carried out at 37°C and 5% CO_2_ concentration in an incubation chamber fitted to a Keyence BZX700 fluorescence microscope. Changes in DiBAC fluorescence were measured within an hour following incubation to ensure that the dye signal reached a steady state at excitation and emission wavelengths of 488 and 520 nm, respectively. Three regions of interest per well and three wells per condition were imaged. DiBAC mean pixel intensity excluding background values was measured using ImageJ.

### Electrophysiology

Patch clamp experiment was conducted in the whole cell configuration at 37±1 C. Cells were passaged and 1.5 × 10^5^ cells were seeded in an uncoated 2cm plastic dish 24 hours prior to the experiment. On the day of the experiment cells were superfused with an extracellular solution containing (in mM): NaCl 140, KCl 5.4, CaCl_2_ 1.8, MgCl_2_ 1, HEPES-NaOH 10, Glucose 5.5, pH=7.4. 1mM of BaCl_2_ and 10 mM of 4-AP (4-Aminopyridine) were added to the extracellular solution to dissect Kir2.1-mediated and Kv1.5-mediated currents, respectively. Either RFP- or GFP-positive cells were identified and used for the analysis. Borosilicate glass pipettes had a resistance of 5-7 MΩ when filled with an intracellular-like solution containing (in mM): 130 K-Gluconate, 10 NaCl, 5 EGTA-KOH, 2 MgCl_2_, 2 CaCl_2_, 2 ATP (Na-salt), 5 creatine phosphate, 0.1 GTP, 10 HEPES-KOH; pH 7.2. Resting membrane potential (RMP) recordings were carried out in the I/0 configuration; after the stabilization of the signal for more than 10 seconds, the extracellular solutions containing either treatments (BaCl_2_ and 4-AP) were exchanged until stabilization of the RMP. Kir2.1-mediated current was recorded in the voltage clamp configuration with a ramp protocol from −100 mV to 100 mV (duration of 100 ms) preceded by a 20 ms long step at −100 mV from a holding potential of 0 mV and analyzed as the Barium-sensitive current. Kv1.5-mediated current was recorded in the voltage clamp configuration with voltage test steps from a holding potential of 0 mV in the range −60/80 mV (ΔV=+10 mV; duration of 1.75 s each step) and analyzed as 4-AP-sensitive current. Currents were normalized to cell capacitance. Density current-voltage plots were then reconstructed for both conductances; for the Kv1.5-mediated current, steady state current was used. Cell capacitance compensation was applied during current recordings and liquid junction potential correction was applied as previously reported^63^.

### In vitro drug studies

MDA-MB-231 and MDA-MB-468 cells were seeded at 5 × 10^3^ cells/well on uncoated 96 well polystyrene plates allowed to adhere for 24 hours. Cells were then treated with various concentrations (0.001, 0.01, 0.1, 1, 5, 10, 50, 100, 500, and 1000 μM) of Amiodarone hydrochloride (40-955-0; Tocris Bioscience, Minneapolis, MN), Carvedilol (C3993,Sigma, St. Louis, MO), Imipramine hydrochloride (I7379; Sigma, St. Louis, MO), and Thioridazine hydrochloride (T9025, Sigma, St. Louis, MO) reconstituted in DMSO according to the manufacturer’s instructions, or 1% DMSO and incubated for 3 days. After establishing IC50 values, 5 and 10μM was chosen for downstream assays.

### Immunocytochemistry

Cells were plated on 96 well polystyrene dishes (MatTek, Ashland, MA) and allowed to adhere for 24 hours. Cells were then fixed for 20 min in 4% paraformaldehyde, then permeabilized with 0.2% TritonX-100, blocked with 5% BSA and incubated with primary antibodies overnight at 4°C. Cells were stained using DAPI (D1306; ThermoFisher Scientific, Waltham, MA) and phalloidin (A12390; ThermoFisher Scientific, Waltham, MA) and simultaneously incubated with fluorescently labeled secondary antibodies at room temperature for one hour. Imaging was performed using a Keyence BZ-X710 microscope (Keyence, Elmwood park, NJ) and mean fluorescent intensity was quantified using ImageJ.

### Western blot

Standard procedures were used for protein electrophoresis and Western blotting. Protein lysates were collected from 231-Control, 231-Kv1.5, 231-Kir2.1 with and without *CDH11* siRNA treatment and separated by SDS–polyacrylamide gel electrophoresis, transferred to a nitrocellulose membrane, blocked with 5% nonfat dry milk solution, and incubated in primary antibody overnight at 4°C. Proteins were detected using horseradish peroxidase–conjugated secondary antibodies. Imaging was performed using a ChemiDoc MP imaging system (12003154; Bio-Rad, Hercules, CA).

### Cell viability assay

Cells were seeded on uncoated polystyrene plates allowed to adhere for 24 hours. Medium was then changed to medium containing vehicle (DMSO) or ion channel drug and incubated for 48 hours. PrestoBlue^TM^ Cell Viability Reagent (A13261; Invitrogen, Carlsbad, CA) was added to each well according to the manufacturer’s instructions and incubated for 25 min at 37°C. Fluorescence was then read on a plate reader at 562nm. Background was corrected to control wells containing only cell culture media (no cells). A dose-response curve was generated for each drug and the IC50 was calculated. Data are the result of at least three independent experiments with three technical replicates per experiment.

### 2D migration assay

For 2D migration, cells were plated on uncoated plastic dishes (MatTek, Ashland, MA) and allowed to adhere for a minimum of one hour. Cells were imaged overnight with images acquired every 10 min for 16 hours in an environmentally controlled chamber within the Keyence BZ-X710 microscope (Keyence, Elmwood park, NJ). Cells were then tracked using VW-9000 Video Editing/Analysis Software (Keyence, Elmwood Park, NJ) and both cell speed and distance migrated were calculated using a custom MATLAB script vR2020a (MathWorks, Natick, MA). Data are the result of at least three independent experiments with three fields of view per experiment and an average of eight cells tracked per field of view.

### 3D invasion assay

Cells were seeded in low-attachment plates in media, followed by centrifugation to form spheroids. Spheroids were grown for three days after which matrix was added to each well, which includes collagen I protein, 10mM NaOH, 7.5% 10x DMEM and 50% 1x DMEM. The spheroids in matrix were then spun down and a further 50μL of media added to each well. Following another seven days of growth (three days for amiodarone treatment studies), spheroids were imaged as a Z-stack using a Keyence BZ-X710 microscope (Keyence, Elmwood Park, NJ) and Z-projection images analyzed using ImageJ. Data are the result of at least three independent experiments with three technical replicates per experiment.

### Cell adhesion assay

Cells were plated on glass-bottomed dishes (MatTek, Ashland, MA) coated with 20μg/ml collagen I and allowed to adhere for 24 hours. Cells were then fixed for 20 min in 4% paraformaldehyde, then permeabilized with 0.2% TritonX-100, blocked with 5% BSA and incubated with primary antibodies overnight at 4°C. Cells were stained using DAPI (D1306; ThermoFisher Scientific, Waltham, MA) and phalloidin (A12390; ThermoFisher Scientific, Waltham, MA) and simultaneously incubated with fluorescently labeled secondary antibodies at room temperature for one hour. Imaging was performed using a Keyence BZ-X710 microscope (Keyence, Elmwood park, NJ) and CellProfiler v3.1.8 (Carpenter et al) was used for imaging analysis using a custom pipeline to quantify cell area and shape parameters based on the phalloidin channel.

### RNA sequencing and analysis

RNA was extracted from three separate biological replicates of MDA-MB-231-Control, 231-Kv1.5, and 231-Kir2.1 cells using the the RNA MiniPrep Kit (R1057; Zymo Research; Irvine, CA). Samples were confirmed to have sufficient quality with Agilent Bioanalyzer before undergoing library preparation by Tufts University Genomics Core using the TruSeq stranded mRNA kit (20020594; Illumina; San Diego, CA). Samples were run on a HiSeq2500 platform at a depth of 50 million single-end 50 base pair reads to allow for transcript quantitation^64^. Quality control of raw read was performed using FastQC^65^ and adapter trimming and read filtering was done using Trimmomatic^66^ with standard settings. Reads were mapped to the reference human genome (assembly GRCh38) with STAR^67^. Differential gene expression analysis was performed using EdgeR in R^68,69^, genes with cpm less than 1 were filtered, and a fold change threshold of 1.5 was used. Pathway enrichment analysis was performed using Gen Set Enrichment Analysis (GSEA)^70,71^. Raw reads and processed data files were uploaded to the Gene Expression Omnibus (GEO) under accession code GSE171150.

### *CDH11* knockdown

Knockdown of *CDH11* was conducted as described previously^53^. MDA-MB-231 cells were passaged and incubated at 1.0 × 10^5^ cells per well in a six well polystyrene coated plate for 24 hours. Cells were transfected with On-Target^TM^ plus Human *CDH11* siRNA-Smartpool (L-013493-00-0005, Dharmacon, Lafayette, CO, US) that contains 4 siRNA mix. *CDH11* #1: 5’-GUGAGAACAUCAUUACUUA-3’. *CDH11* #2: 5’- GGACAUGGGUGGACACAUAUG-3’. *CDH11* #3: 5’-GGAAAUAGCGCCAAGUUAG-3. *CDH11* #4: 5’-CCUUAUGACUCCAUUCAAA-3’ using Lipofectamine RNAiMAX transfection reagent (13778030; ThermoFisher Scientific, Waltham, MA) in serum-free DMEM. Silencer^TM^ Negative Control #1 siRNA (AM4611; ThermoFisher Scientific, Waltham, MA) was used as a negative control for transfection. The final concentration of siRNA was 80nM. The siRNA-transfected cells were incubated for 48 h and used for functional assays.

### Animal experiments

Animal procedures were carried out in full accordance with established standards set forth in the Guide for the Care and Use of Laboratory Animals, 8th edition (NIH Publication No. 85-23). For xenograft tumors, 2 × 10^6^ MDA-MB-231 cells in PBS and 20% collagen I were injected into the fourth right mammary fat pad of 6-wk-old female NOD-SCID-γ (NSG) mice (005557; JAX, Bar Harbor, ME). For the channel overexpressing lines, 10 mice each were injected with either MDA-MB-231 control, Kir2.1, or Kv1.5 cells and tumor growth was monitored for seven weeks. For the amiodarone experiment MDA-MB-231 GFP cells were injected into 20 animals and 0.05mg/kg amiodarone or DMSO control was administered i.p. beginning at five weeks post-injection every day for two weeks, followed by injection of 0.5mg/kg amiodarone or DMSO control for an additional 10 days. During the drug injection period, tumor growth was measured biweekly and body weight weekly. Mice were euthanized by CO2 and tumors and lungs were excised once tumors reached a volume of 1.5cm^3^ or endpoint were met.

### Histology

Fixation, processing and staining of tissue sections from tumors and lungs was carried out as previously described^72^. Tumors and lungs dissected from NSG mice were fixed in 10% paraformaldehyde and embedded in paraffin. Tissue was sectioned using a microtome at 10μm thickness. For H&E staining: standard procedures were followed, including deparaffinizing, hydration, staining with Hematoxylin (GHS280; SIGMA, St. Louis, MO) and counterstaining with Eosin (HT110180; SIGMA, St. Louis, MO).

### Statistical analysis

GraphPad Prism v8.4.3 was used for generation of graphs and statistical analysis. For data with normal distribution: for comparison between two groups, an unpaired two-tailed Student’s t-test was used and a p-value of ≤0.05 considered significant, and for comparison between multiple groups a one-way ANOVA with Dunnett’s multiple testing correction was used with a corrected p-value of ≤0.05 considered significant. Data represent mean ±S.E.M. unless otherwise stated. Patch clamp data were analyzed with Clampfit10 (Axon) and Origin Pro 9. Linear fitting was performed reporting the adjusted R^2^ value.

## Data Availability

All RNA-seq raw data generated for the present study, along with counts matrices and metadata for each sample, are publicly available in GEO under accession code GSE171150. All other data supporting the findings of this study are available from the corresponding author upon reasonable request.

## Acknowledgements

We thank Joan Lemire for plasmid preparation, Mattia Bonzanni for performing electrophysiology experiments, and David Kaplan for support. We also thank members of the Oudin lab for manuscript review. We thank Allen Parmelee and Steven Kwok at the Tufts University School of Medicine Laser Cytometry core facility for performing FACS. We also thank the Tufts University Core Facility Genomics for performing RNA sequencing.

## Author Contributions

S.L.P, M.L., and M.J.O. conceived the project, designed the study, and interpreted results. S.L.P. derived ion channel overexpressing cell lines used in the study and performed DiBAC, 2D migration, 3D invasion, immunostaining, western blot, imaging, and siRNA knockdown assays. S.L.P., P.R., and D.H.S. performed image analysis for immunostaining and the 3D invasion assay. D.H.S. performed cell area and morphology analysis. S.L.P. and T.T.L. performed RNAseq analysis. P.R. performed in vitro drug studies, generated standard dose curves and analyzed cell viability. S.L.P. and P.R. conducted and analyzed animal drug study. S.L.P. and D.H.S. processed and stained tissue. S.L.P., T.T.L. and P.R. quantified lung tissue. M.L., and M.J.O. provided experimental and analytical support. S.L.P. and M.J.O. wrote the manuscript with feedback from all authors.

## Supplemental Information

**Figure S1:**
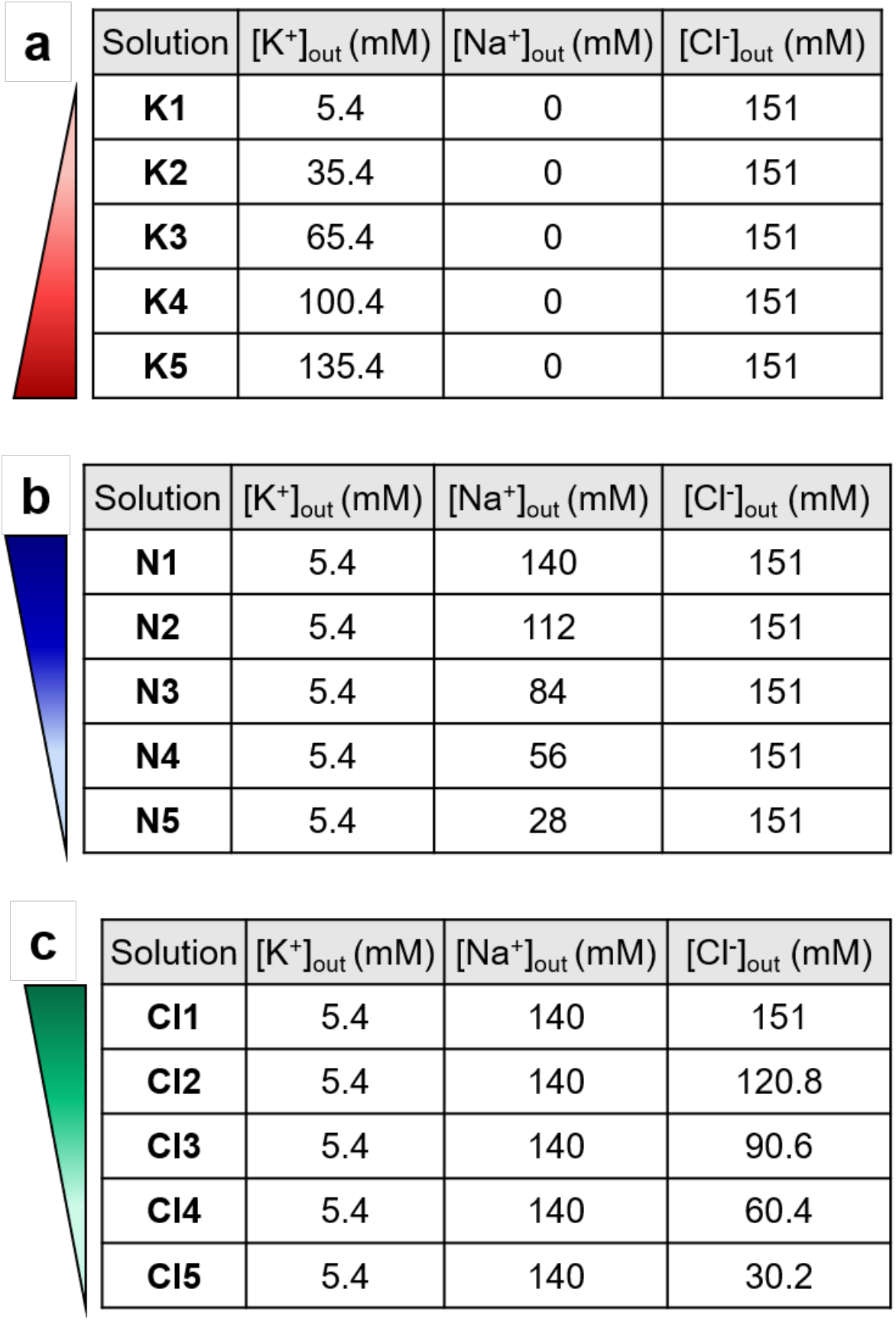
Recipes for ionic solutions to isolate single ion effect on breast cancer cell RMP. Solutions of increasing concentration of one ion species were made as previously described^25^ for K^+^ **(a)**, Na^+^ **(b)**, and Cl^−^ **(c)**. These solutions were osmotically balanced and able to support cell survival for the duration of the assay.

**Figure S2:**
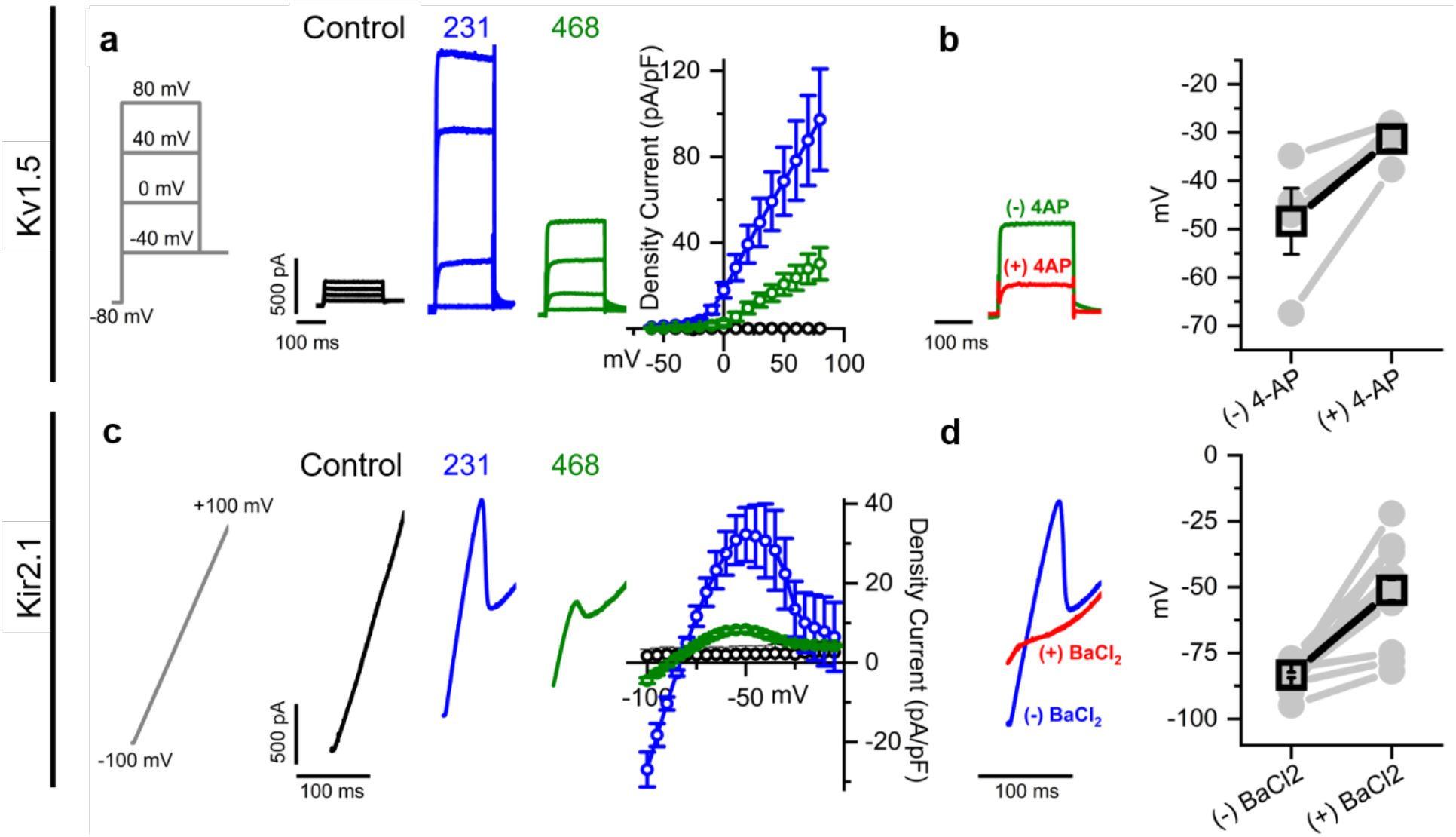
Electrophysiological characterization of stable lines overexpressing either Kv1.5 or Kir2.1 channel. **(a)** Kv1.5-dependent currents (IKv1.5) were assessed applying a depolarizing (ΔV=10 mV) voltage steps from a holding potential of −80 mV (protocol in grey). Representative current traces and the density current-voltage plot are showed for the indicated conditions and voltages (GFP: black, n=6; 231 Line: blue, n=9; 468 Line: green, n=4). **(b)** The identity of the IKv1.5 was confirmed by applying the blocker 4-AP (10 mM). Representative IKv1.5 traces with and without the blocker (red and green traces, respectively). The resting membrane potential (RMP) values are shown before and after application of 10 mM of 4-AP (grey circles); the mean ± S.E.M. Mean RMP values are indicated with black squares. **(c)** Kir2.1-dependent currents (IKir2.1) were assessed applying a ramp protocol in the range −100 mV/100 mV range (duration 100 ms; protocol in grey). Representative current traces and density current-voltage plot are shown for the indicated conditions (GFP: black, n=4; 231 Line: blue, n=11; 468 Line: green, n=11). **(d)** The identity of the IKir2.1 was confirmed applying the blocker BaCl_2_ (1 mM). Representative IKir2.1 traces with and without the blocker (red and blue traces, respectively). The RMP values are shown before and after application of 1 mM of BaCl_2_ (grey circles); the mean ± S.E.M. Average RMP values are indicated with black squares.

**Figure S3:**
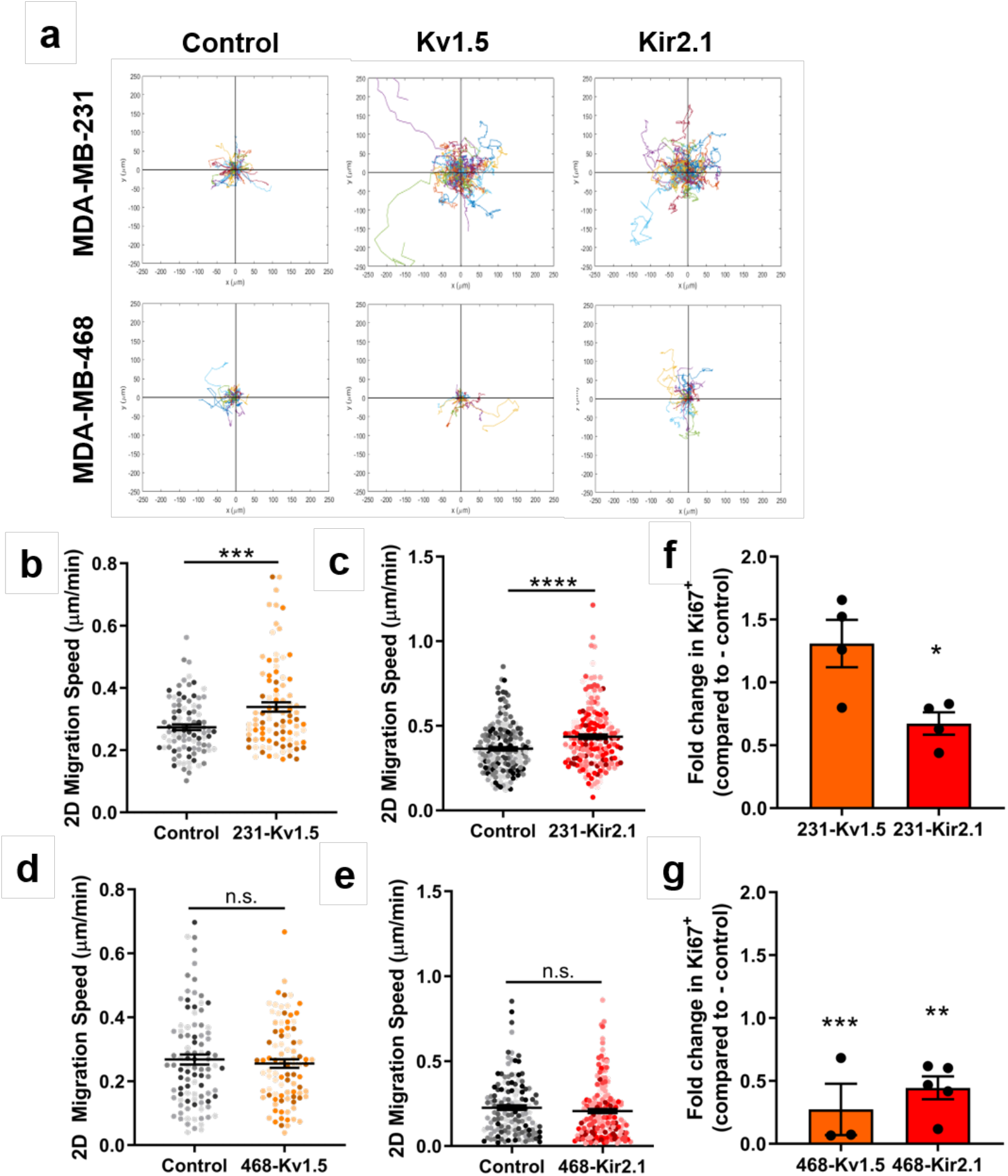
2D migration and proliferation is altered with K^+^ channel-driven hyperpolarization. **(a)** Representative rose plots of tracked 2D migration of 231 and 468 lines overexpressing Kv1.5 and Kir2.1. **(b-c)** 2D migration of 231 lines compared to control (Kv1.5: ***p=0.0002, Kir2.1: ****p<0.0001). **(d-e)** 2D migration of 468 lines compared to control. Fold change in Ki67 expression for **(f)** 231 (*p=0.0104) and **(g)** 468 (**p=0.0081, ***p=0.0010) lines compared to control cells. Data are pooled from three or more biological replicates and shown as mean ± S.E.M. Different color shades within group represent samples from different replicates. Significance was determined using an unpaired two-tailed Student’s t-test.

**Figure S4:**
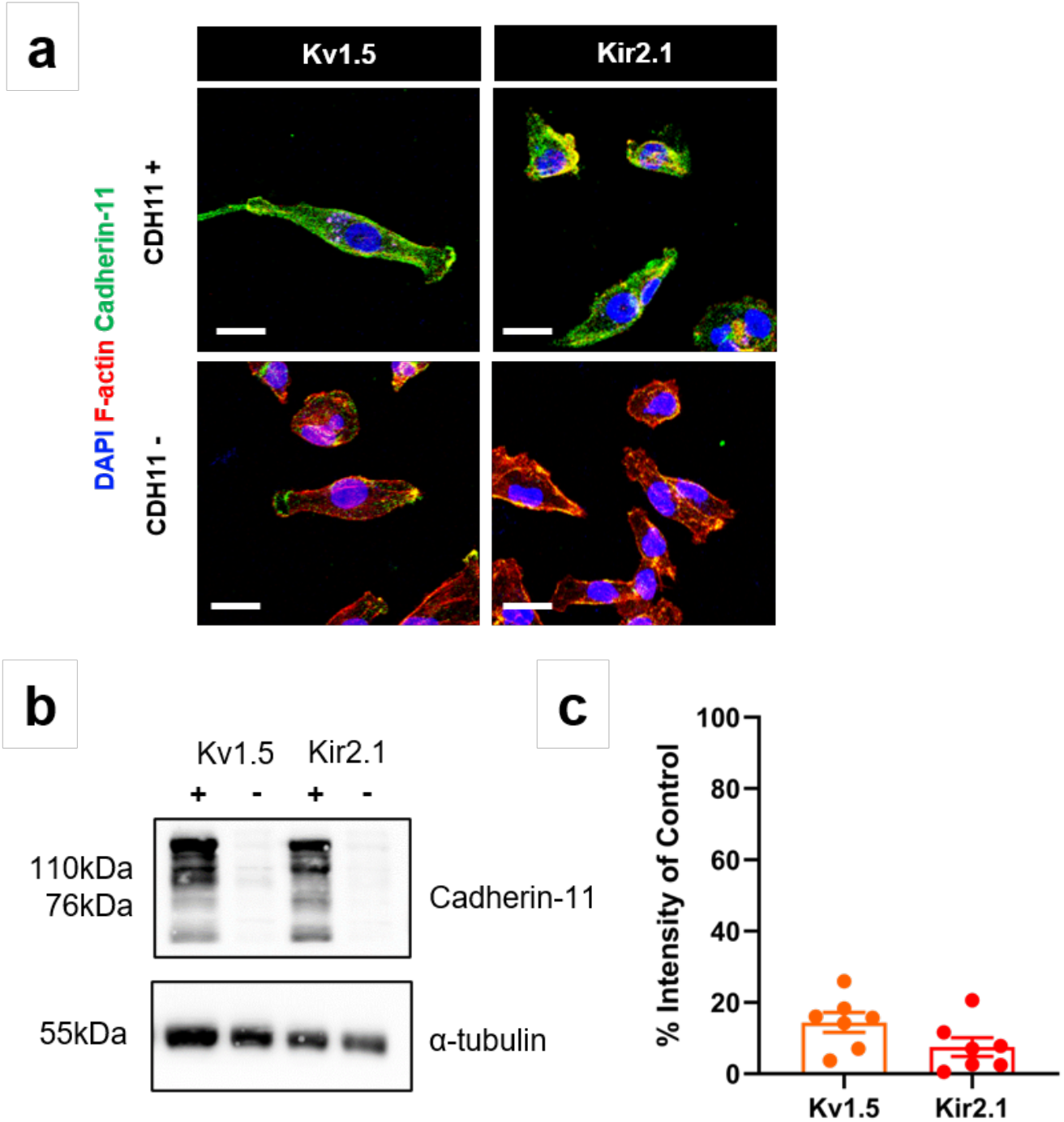
*CDH11* siRNA-mediated knockdown in 231-Kv1.5 and 231-Kir2.1 cells. **(a)** Representative images of cells immunostained for cadherin-11 and counterstained for F-actin. Scale bar = 10μm. **(b)** Representative western blot and **(c)** quantification of cadherin-11 expression with and without *CDH11* siRNA knockdown after 48 hours. Data are shown as mean ± S.E.M., n=6.

**Figure S5:**
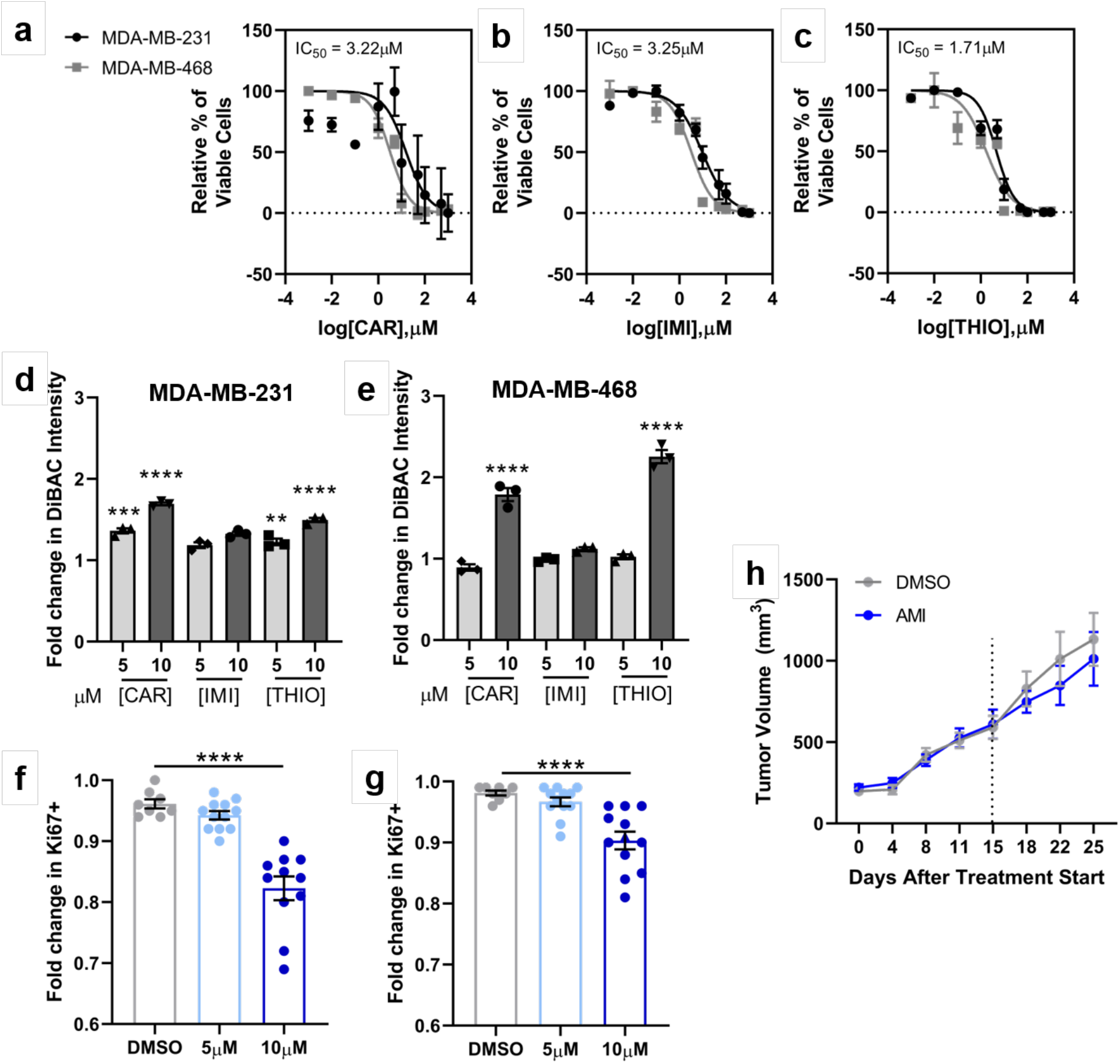
Effect of K^+^ channel blockers on TNBC viability and RMP. Dose response curves of cell viability in MDA-MB-231 and MDA-MB-468 cells cultured with varying concentrations of **(a)** carvedilol (CAR), **(b)** imipramine (IMI), or **(c)** thioridazine (THIO). Quantification of RMP in **(d)** MDA-MB-231 and **(e)** MDA-MB-468 cells treated with carvedilol (CAR), imipramine (IMI) or thioridazine (THIO) as measured by mean DiBAC intensity. Data are expressed as mean ± S.E.M, n=3 biological replicates. Significance was determined using a two-way ANOVA with Dunnett’s multiple comparisons test. **(f)** MDA-MB-231 and **(g)** MDA-MB-468 cell fold change in proliferation after treatment with amiodarone as measured by number of DAPI^+^/Ki67^+^ cells. **(h)** Tumor volume measured over time in animals injected with MDA-MB-231 cells and treated with amiodarone or DMSO. Data shown as mean ± S.E.M. N = 9-10 animals per group.

